# HS3ST1 regulates pulmonary inflammation and is a determinant of clinical outcomes after trauma and hemorrhagic shock

**DOI:** 10.64898/2026.05.07.723543

**Authors:** Ava K. Mokhtari, Madeline E. Cotton, Kimberly A. Thomas, Alisha Chitrakar, Joseph D. Krocker, Marissa Pokharel, Baron K. Osborn, Maria del Pilar Huby Vidaurre, Atharwa R. Mankame, Charles E. Wade, Yao-Wei Wang, David J. Orlicky, Mitchell J. Cohen, Jillian R. Richter, Nicholas W. Shworak, Jessica C. Cardenas

## Abstract

Mechanisms that promote organ injury after trauma and hemorrhagic shock (T/HS) remain poorly defined. Endothelial heparan sulfates with a 3-O-sulfate (3-OS) modification, controlled by the HS3ST1 gene, have anticoagulant and anti-inflammatory properties through their interaction with antithrombin. Our objective was to determine whether HS3ST1 deficiency drives organ injury and poor outcomes after T/HS. *Hs3st1*^-/-^ and wild-type (WT) mice were subjected to T/HS followed by resuscitation with lactated ringer’s (LR) or fresh frozen plasma (FFP). While no differences were observed between WT and *Hs3st1*^-/-^ LR resuscitated mice, lung injury and leukocyte infiltrates were significantly increased in FFP resuscitated *Hs3st1*^-/-^compared to WT mice. In vitro, leukocyte slow rolling and adherence was increased in HS3ST1 KO compared to WT cells. Among 472 T/HS patients, of which 31 (7%) were homozygous for the rs16881446 variant allele (GG), the number of ventilator free days was lower, and mortality was significantly higher in AG and GG patients. The rs16881446 genotype was independently associated with mortality. In conclusion, HS3ST1 deficiency mitigates organ protection from FFP resuscitation, partly through mediating EC:leukocyte engagement, and predicts mortality after T/HS. These findings identify a novel therapeutic target and prognostic tool that can be leveraged towards improved risk stratification after trauma.

## Introduction

Organ injury and subsequent development of multiple organ dysfunction syndrome (MODS) remain key contributors to late-stage morbidity and mortality after traumatic injury and hemorrhagic shock (HS).(1, 2) Our current mechanistic understanding of post-trauma organ injury stipulates a key role for thromboinflammation, or concurrent dysregulation of the coagulation and inflammatory systems that facilitate end-organ leukocyte infiltration, perivascular fibrin deposition, and edema; however, the precise cellular and molecular pathways that promote thromboinflammation remain poorly resolved.(2, 3) Consequently, there are no therapeutic countermeasures available to target thromboinflammation and prevent or treat late-stage MODS.

Microvascular endothelial cells (EC) regulate coagulation, inflammation, and vascular barrier function, in part, through their expression of cell surface glycosaminoglycans, of which heparan sulfates are the most abundant.(4, 5) Heparan sulfates are structurally heterogenous, with their function and interactions with blood proteins determined by sulfation motifs added post-translationally to the disaccharide backbone.(6, 7) Heparan sulfates with in vitro anticoagulant activity are uniquely defined by their ability to bind and activate antithrombin (AT), which is dependent on sulfation at the 3-*O* position of the central glucosamine, a modification that is controlled almost exclusively by the EC 3-*O*-sulfotransferase enzyme encoded by the HS3ST1 gene.(8–10) Similar to heparin, the interaction between AT and EC 3-*O*-sulfated (3-OS) heparan sulfate accelerates AT-mediated inhibition of factor Xa (FXa) and thrombin in vitro, but such anticoagulant activity does not appear to be operable in vivo.^13^ However, AT bound to 3-OS heparan sulfate also induces intracellular prostacyclin synthesis, which attenuates platelet and leukocyte activation and inhibits nuclear translocation of NFκB, thereby suppressing EC pro-thromboinflammatory gene expression. (11, 12)

Prior work has demonstrated that deletion of *Hs3st1* in mice significantly reduces EC expression of 3-OS HS, abolishing AT binding to the vessel wall by 90%, resulting in accelerated lethality and increased systemic inflammation following septic challenge.(9, 13) In humans, a common single nucleotide polymorphism (SNP) in the transcriptional regulatory region of the HS3ST1 gene (rs16881446) was associated with development of more severe cardiovascular disease in chronically ill patients(9); however, the role of HS3ST1 and 3-OS heparan sulfate in regulating thromboinflammation and organ injury following trauma and HS remains unexplored. Our previous data show that 1) AT is an important organ protective constituent of fresh frozen plasma (FFP)(14), 2) trauma and HS in rats reduces pulmonary 3-OS heparan sulfate expression, and 3) pharmacologically blocking AT-EC interactions with the heparan sulfate antagonist, surfen, resulted in worse lung injury in FFP resuscitated mice,(15) highlighting the potential importance of 3-OS heparan sulfate expression in development of MODS and ultimately treatment responses in trauma patients. The goal of this study was to determine whether HS3ST1 deficiency drives thromboinflammation, organ injury, and poor outcomes following trauma and HS. We hypothesized that *Hs3st1* deficient mice would be resistant to the organ protective effects of FFP and exhibit worse organ injury compared to wild-type (WT) controls, with an emphasis on the role of 3-OS heparan sulfate in limiting leukocyte engagement to endothelium. We further hypothesized that trauma patients who are homozygous for the variant allele of the rs16881446 SNP would have increased complications and mortality compared to those expressing WT alleles.

## Materials and Methods

### Animal Model of Trauma and Hemorrhagic Shock

Animal procedures were conducted under a protocol approved and strictly followed by the University of Colorado (01398) and in accordance with Animal Research: Reporting of In Vivo Experiments (ARRIVE) guidelines. *Hs3st1*^+/+^ (wild type; WT) and *Hs3st1*^-/-^ (knock out, KO) mice were generated on an isocongenic C57BL/6J background, as previously described.(13) Prior work has confirmed by mass spectrometry that *Hs3st1*^-/-^ mice express minimal 3-OS heparan sulfate.(16) Male and female mice (10-14 weeks old) were use for experimentation. A well-described model of laparotomy followed by fixed pressure hemorrhage was used to induce soft tissue trauma and HS, as we have used previously.(14, 15, 17) Briefly, mice were placed in a supine position on a temperature-controlled surface and maintained under isoflurane anesthesia for the duration of the experiment. The right femoral artery and vein were cannulated with heparin-washed PE-50 polyethylene catheters and the arterial blood pressure, mean arterial pressure (MAP), and heart rate were monitored and recorded continuously (PowerLab4/10 and LabChart Pro, AD Instruments). Following a 15-minute stabilization period, a baseline blood sample was collected for blood gas measurements (iSTAT, McKesson). A midline laparotomy (3 cm) exposed the bowels which were exteriorized with gentle distention of the small intestine to induce soft tissue trauma, followed by repositioning of the small intestine and laparotomy closure. HS was induced by rapid, controlled hemorrhage from the arterial catheter to a target MAP of 35 mmHg, which was sustained for approximately 1 hour through fluid infusion and removal until decompensation occurred. At this time, another blood sample was collected for blood gas measurements and resuscitation was initiated to restore baseline MAP with either crystalloid (lactated ringers; LR) or mouse pooled fresh frozen plasma (FFP) based on randomization group. Mice were awakened from anesthesia and monitored for 3 hours. After the observation period, mice were re-anesthetized and a final blood sample was collected, lungs were extracted, flash frozen in liquid nitrogen with a portion of lung tissue fixed in 10% formalin, and the animals were humanely euthanized. Sham animals underwent anesthesia, vessel cannulation, and observation with no laparotomy or HS prior to tissue collection.

### Tissue Analysis

Lungs were fixed in 10% formalin, paraffin embedded, sectioned, and stained with hematoxylin and eosin (H&E). Hematoxylin and eosin (H&E) stained lung tissue was analyzed by an independent, blinded histopathologist for histologic lung injury at 40x objective, expressed as arbitrary units. Tissues were scored using a 4 point scale (**Supplementary Figure 1**). The total lung injury score was calculated by multiplying the scores from each category by the score for percentage of affected tissue. Lung tissue was stained with an anti-myeloperoxidase (MPO) antibody (Abcam ab208670; 1:750) to quantify neutrophil infiltrates using 3 random images of each lung. The number of MPO-positive cells per field of view was counted by an individual blinded to the treatment groups. The presence of intravascular fibrin deposition was evaluated using Carstairs staining (StatLab, McKinney, TX, USA). Three images containing pulmonary capillaries were collected for each lung and ImageJ was used to quantify fibrin-positive (red) pixels and expressed as percent positive staining normalized to each vessel circumference. Total RNA was extracted from frozen lung tissue of WT sham, WT FFP, KO sham and KO FFP mice using an RNAspin mini RNA isolation kit (25-0500-72, Cytiva, Marlborough, MA, United States) according to manufacturer’s protocol. Gene expression was assessed by bulk RNAseq (Novogene, Ashland, MA). Transcriptomic analysis was performed using RStudio in the R environment (version 2025.5.1+513). Differentially expressed genes (DEGs) were determined based on statistical thresholds of -log10 p-value > 1.3 and an absolute log2 fold-change > 0.5. *EnhancedVolcano* and *pheatmap* packages(18) were used to generate volcano plots and heatmaps, respectively. The STRING database (v12.0) was used for unbiased functional enrichment analysis of DEGs associated with WT and KO responses to trauma, HS and FFP resuscitation. Genes were stratified based on the direction of the genotype x injury interaction effect, and Reactome pathways were identified based on ≥10 overlapping genes and false-discovery rate (FDR)-adjusted p < 0.05.

### In vitro Perfusion Assays

EA.hy926 cells (a hybrid fusion of human umbilical veinous endothelial cells and A549 cells [adenocarcinomic human alveolar basal epithelial cells]) were used as an endothelial cell surrogate. HS3ST1 deletion was performed using CRISPR, allowing for the creation of EA.hy926 HS3ST1^-/-^ (“KO”) cells. The Cas9 and single guide RNA were delivered via lentiviral transfection. WT and KO clones were identified by sequencing. HS3ST1^+/+^ parental EA.hy926 cells were used as “WT”. Cells were cultured in complete media (Vascular Cell Basal Medium (#PCS-100-030, ATCC, Manassas, VA, USA), with Endothelial Cell Growth Kit-VEGF (#PCS-100-041, ATCC), and 100 units/mL penicillin/streptomycin (#15-140-122, Fisher Scientific, Waltham, MA, USA)) in fibronectin-coated (#1030-FN-05M, R&D Systems, Minneapolis, MN, USA) tissue culture treated flasks, and maintained at 37°C and 5% CO_2_. For perfusion assays, WT and KO cells were seeded into fibronectin-coated ibi-Treated µ-slides (VI 0.4, #80606, ibidi, Gräfelfing, Germany) the day before perfusion experiments at 75,000 cells per channel in complete media. Subsequent stimulations and leukocyte perfusions were performed under flow settings of 0.5 dynes/cm^2^ using the ibidi pump, fluidic unit, and white perfusion set (length 50 cm, ID 0.8 mm, 10 ml reservoirs, #10963) at 37°C and 5% CO_2_ coupled with an EVOS M7000 Imaging System with the onstage incubator attachment. For TNFα (#10291-TA-050, R&D Systems) stimulation, channel-seeded WT and KO cells were incubated with complete media containing 50 ng/mL TNFα overnight at 37°C and 5% CO_2_ prior to perfusion with leukocytes. For AT (#HCATIII-0120, Prolytix, Essex Junction, VT, USA) stimulation, 30 min prior to TNFα, seeded cells were incubated with 150 µg/mL AT (Prolytix, Essex Junction, VT, USA). TNFα and AT were administered through an in-line injection port in the ibidi perfusion setup.

Polymorphonuclear cells (PMN) were isolated using Polymorphprep (#AXS-1114683, Cosmo Bio USA, Carlsbad, CA, USA; used per manufacturer’s protocol) from ∼20mL whole blood collected into sodium citrate from adult healthy human donors (N=3). Donors were consented and enrolled in accordance with Vitalant Research Institute’s approved IRB protocols, and blood collected per Vitalant’s standard collection protocols. Isolated PMN were washed with PBS and labeled with CFSE (#423801, Biolegend, San Diego, CA, USA) for 20 min at 37°C, then the staining reaction quenched, and cells resuspended to 0.5X10^6^/mL and stored at 37°C until use.

CFSE-stained PMN were perfused over WT or KO EC at 0.5 dynes/cm^2^ to allow for video capture and enumeration of slow rolling and adherent PMN on the surface of stimulated EC. Twenty second videos of GFP signal were captured at 5-7 unique sites within each channel using a 20X objective (**Supplemental Videos 1 & 2**). Videos were analyzed using NIS Elements AR Software (Nikon, Melville, NY, USA) using object identification and tracking features. Detailed NIS Elements analysis procedures are available upon request. In brief, GFP+ (CFSE-labeled) objects were counted, and the speed of each object reported as pixels/sec, which was converted to μm/sec. Speed was binned by 0-50 μm/sec (slow rolling, tethering, and adherent PMN), 50-250 μm/sec (fast rolling PMN), and >250 μm/sec (non-engaging with EC). To enumerate the frequency of PMN at each speed, the following calculation was used: % of PMN = [# of objects @ speed X ÷ total # of objects]*100. With respect to reproducibility, assays were performed on a given day (unique EC passage) with a unique PMN donor (N=3 total), such that each experimental day was a unique biological replicate.

### Human Subjects

Whole blood was collected from patients enrolled in a single center prospective observational study conducted at Memorial Hermann Hospital – Texas Medical Center in Houston, Texas between July 2017 and June 2023. The study was approved by the University of Texas Health Science Center at Houston Institutional Review Board (HSC-GEN-0059) and enrolled all adult (≥ 16 years) highest level trauma activation patients for whom consent was obtained within 72 hours from admission. Patients presenting to the hospital in HS, defined as base excess ≤ -4 mmol/L, who received blood transfusions of packed red blood cells (RBC) or whole blood, and consented for use of genetic material were selected for analysis. Patients who were pregnant, prisoners, or who did not consent were excluded. Patient demographics, injury characteristics, blood transfusion, laboratory values, and outcomes were collected from the medical records. Acute kidney injury was defined as having a KDIGO score ≥2.(19)

### Blood Collection and Analysis

Whole blood was collected upon admission to the emergency department into EDTA vacutainers and centrifuged twice at 1,800 x *g* for 10 minutes. Buffy coat layers and platelet poor plasma were removed and stored at -80°C until analysis. Deoxyribonucleic acid (DNA) was extracted from frozen buffy coats using Gentra Puregene Tissue Kits (Qiagen). Real-time polymerase chain reaction (PCR) utilizing the TaqMan SNP Genotyping Assay (Thermo Fisher Scientific) and a StepOnePlus Real-Time PCR System (Applied Biosystems) were performed for *HS3ST1* (rs16881446) genotyping according to the manufacturer’s instructions. TaqMan Genotyper software (Thermo Fisher Scientific) was used for allele discrimination analysis. Plasma inflammatory mediators were quantified using Meso Scale Discovery V-PLEX Human Biomarker 39-Plex kit (MSD, Rockville, MD). Antithrombin (R&D Systems, Minneapolis, MN) and thrombin-antithrombin complex (Innovative Research, Novi, MI) were measured by enzyme linked immunosorbent assay (ELISA).

### Statistical Analysis

All data were assessed for normality using the Kolmogorov-Smirnov test. Normally distributed data are presented as means with standard deviation whereas data that are not normally distributed are presented as medians with interquartile ranges. For normally distributed data, comparisons between groups were determined using one-way analysis of variance using Tukey’s adjustment for multiple comparisons. For data that was not normally distributed, data were compared using Kruskal Wallis test. For perfusion assays, data are represented as mean ± standard error of the mean, and two-way analysis of variance was used to evaluate PMN speed with respect to EC genotype and stimulation. For comparing dichotomous outcomes in patient cohorts, Chi-square tests were used. Logistic regression analyses were conducted to determine the independent association between rs16881446 genotype and death, while controlling for age, sex, and injury severity scores. Logistic regression analyses were conducted using Stata whereas all other analyses were conducted using Graphpad Prism.

## Results

### Hs3st1 *does not modulate severity of, or recovery from hemorrhagic shock*

WT and *Hs3st1* KO littermates were subjected to trauma and HS, followed by resuscitation with either LR or FFP (**Figure 1A**). Subgroup analyses by sex determined there were no significant differences in hemodynamic or metabolic indicators of shock between males and females, therefore data for both sexes was combined. There were no differences across resuscitation groups or between WT and KO mice in the percentage blood volume withdrawn to induce HS (**Figure 1B**) or in the resuscitation volumes necessary to reverse HS (**Figure 1C**). While the final MAP after resuscitation and observation was significantly lower for all mice that underwent trauma and HS compared to sham animals, there were no differences between WT and KO mice or between resuscitation groups (**Figure 1D**). Similarly, final lactate levels were significantly elevated in LR WT, FFP WT, and FFP KO mice compared to sham mice (**Figure 1E**); however, there were no observed differences across genotype or resuscitation groups.

**Figure 1.**
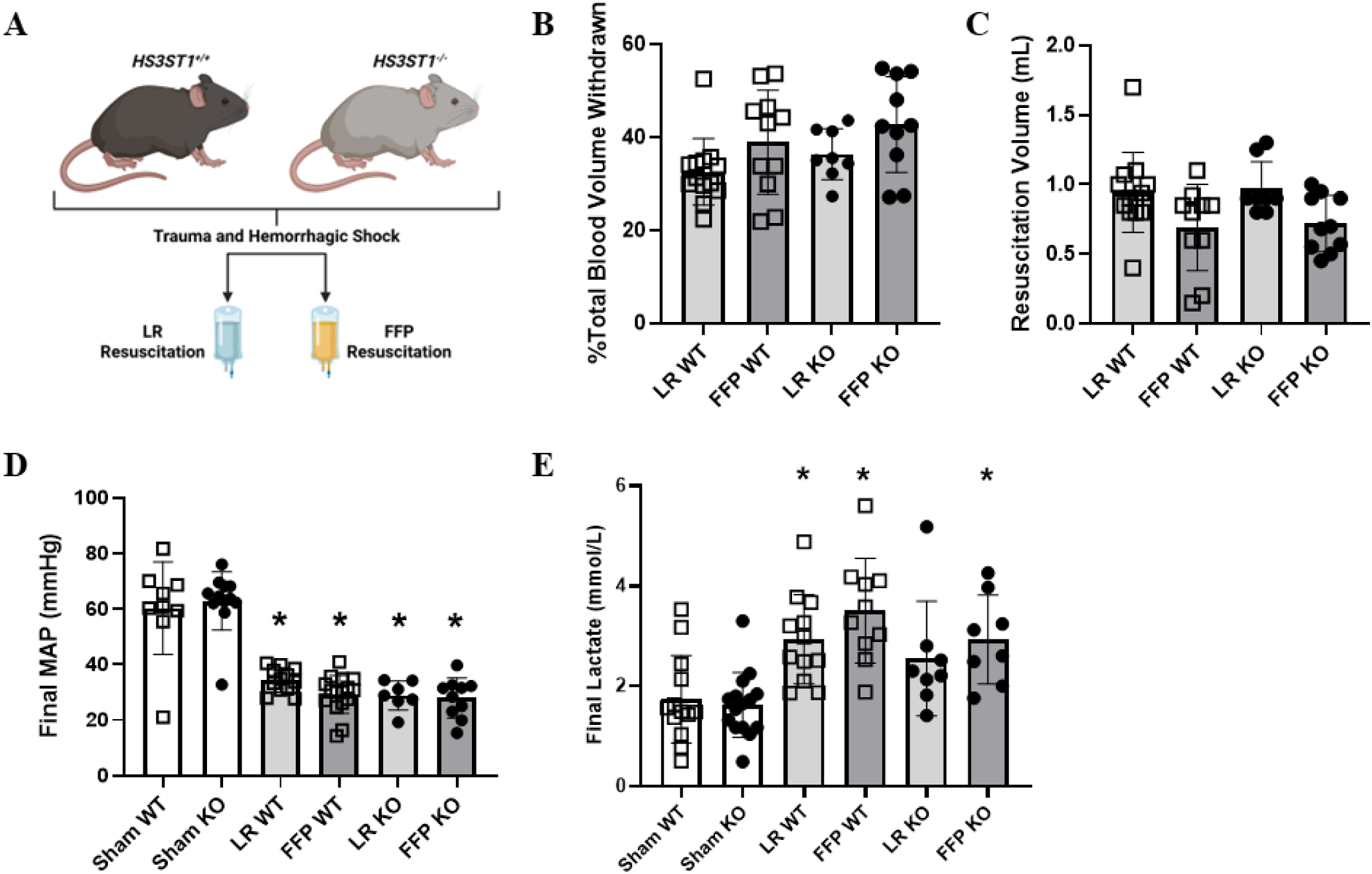
*Hs3st1* deficiency does not modulate the severity of, or recovery from hemorrhagic shock. (A) Schematic depicting experimental schema. The total blood withdrawn to induce HS was expressed as percent of body weight (B). The resuscitation volumes necessary to reverse HS and increase MAP are provided (mL) (C). Final mean arterial pressure (MAP) (D) and final lactate values (E) were recorded at termination. N=8-12 per group. Data reported as mean ± standard deviation. * Denotes significance from sham

### Hs3st1 *deficiency mitigates FFP-mediated reductions in pulmonary neutrophil infiltration after trauma and hemorrhagic shock*

Paraffin embedded lung tissues were stained with Carstairs to determine differences in perivascular fibrin deposition. No differences in fibrin staining were identified across groups (data not shown). Anti-myeloperoxidase (MPO) antibodies were used to quantify the amount of neutrophil infiltration after trauma, HS, and resuscitation. There were no differences in the amount of lung neutrophils between WT and KO sham mice (**Figure 2A**). Compared to WT and KO sham mice, WT mice that underwent trauma and HS and LR resuscitation had greater pulmonary neutrophil infiltrates (70 ± 34 cell/FOV), which was significantly reduced in WT mice resuscitated with FFP (31 ± 12; p<0.01) (**Figure 2A**). Conversely, we found that KO mice resuscitated with LR had significantly increased pulmonary neutrophils (84 ± 38) compared to both sham mice and WT mice resuscitated with LR (both p<0.001). Moreover, KO mice resuscitated with FFP had significant greater pulmonary neutrophil infiltrates compared to sham KO mice (66 ± 26 versus 25 ± 17; p<0.01) and WT mice resuscitated with FFP (p<0.01). There were no significant differences between KO mice resuscitated with LR and KO mice resuscitated with FFP (**Figure 2A**). Representative 20x images are provided (**Figure 2B**).

**Figure 2.**
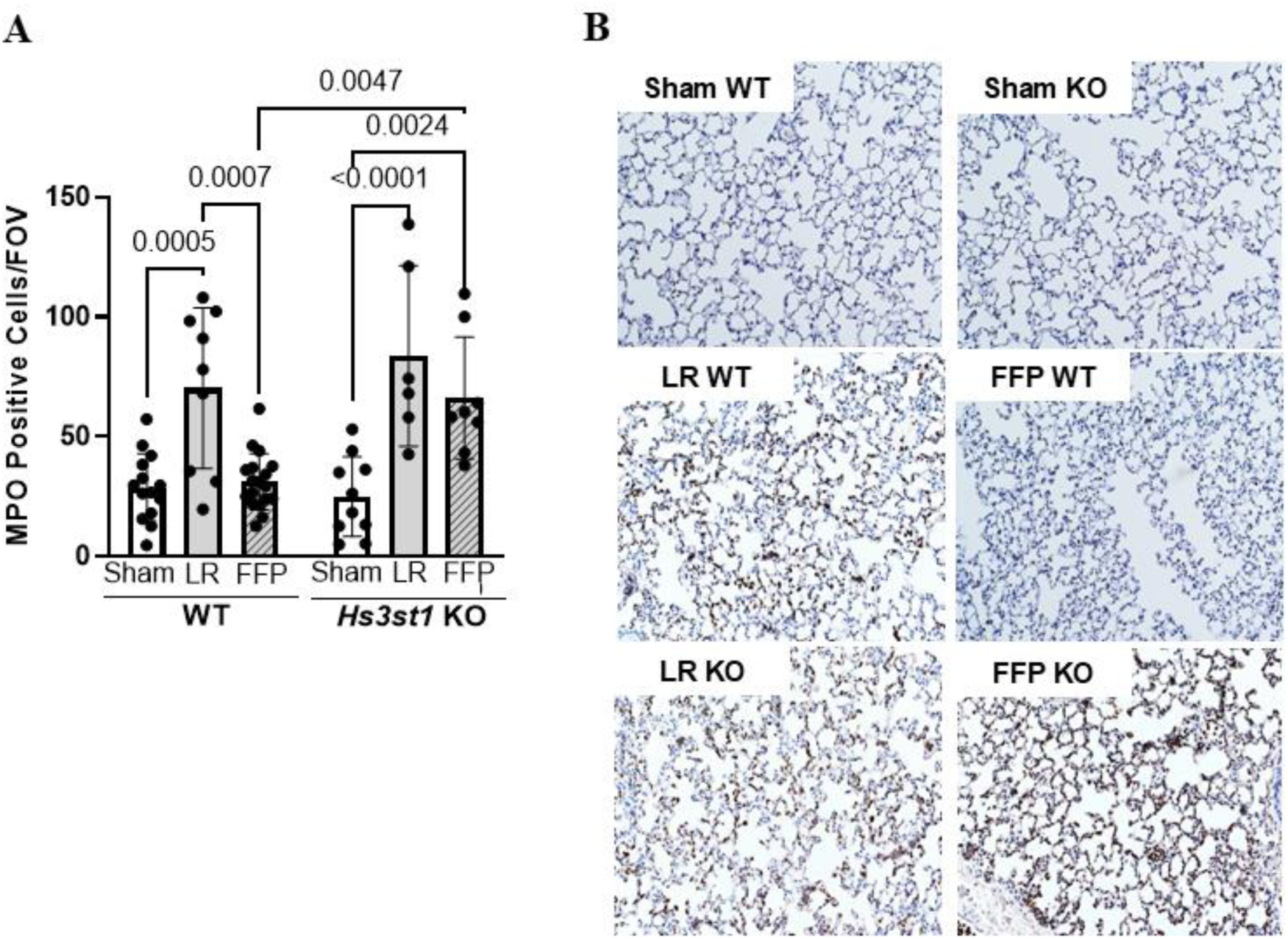
*Hs3st1* KO mice exhibit greater pulmonary inflammation after induction of trauma and HS compared to WT mice. Myeloperoxidase (MPO) positive cells per field of view (FOV) were quantified from 3 random images taken per tissue and averaged for each mouse. Representative images at 20x are provided (bottom). N=6-12 per group. Data reported as mean ± standard deviation.

### Hs3st1^-/-^ mice have more severe histologic lung injury compared to Hs3st1^+/+^ mice after trauma and HS

No differences in histologic lung injury were observed between WT and KO sham mice (**Figure 3A**). Compared to WT and KO sham mice, those that underwent trauma, HS, and LR resuscitation had significantly worse lung injury, which was significantly reduced in WT mice resuscitated with FFP (LR WT 45 ± 12 arbitrary units versus FFP WT 26 ± 5.7 arbitrary units; p=0.01). KO mice resuscitated with LR had significantly increased lung histology scores compared to all sham mice and WT mice resuscitated with FFP; however, lung injury scores were similar between WT and KO mice resuscitated with LR. When examining KO mice resuscitated with FFP, we found that lung injury scores (50 ± 15) were significantly elevated compared to sham and WT mice resuscitated with FFP (p<0.01) and there was no difference when comparing KO mice resuscitated with FFP to those resuscitated with LR (**Figure 3A**). Representative 20x images are provided (**Figure 3B**).

**Figure 3.**
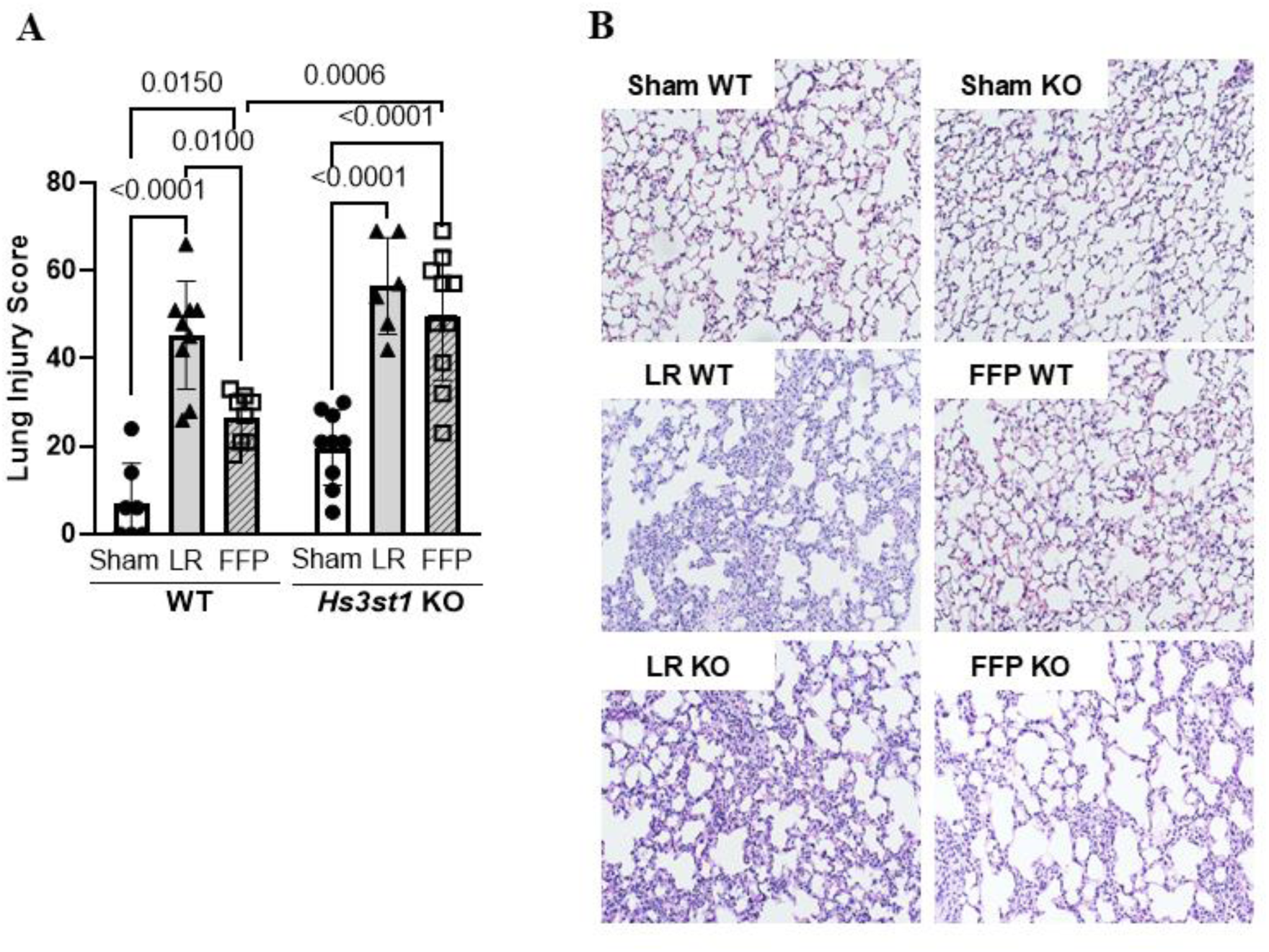
Histologic lung injury is more severe in *Hs3st1* KO mice compared to WT after trauma and HS. H&E stained lung slices were evaluated by a blinded histopathologist and scored for histologic injury. Representative images at 20x are provided (bottom). N=6-12 per group. Data reported as mean ± standard deviation.

### Hs3st1^-/-^ *mice exhibit altered transcriptional responses to trauma, HS and FFP resuscitation compared to WT mice*

Given that FFP protection against post-injury pulmonary inflammation was observed in WT but not Hs3st1 KO mice, exploratory RNA sequencing analysis was performed on total lung tissue to identify transcriptional differences in WT and KO responses to provide insight into the role of 3-OS heparan sulfate in regulating organ damage. In WT mice, a total of 3387 DEG were identified between sham and trauma/HS with FFP resuscitation (**Figure 4A**), and 2038 DEG were observed between KO sham and KO FFP mice (**Figure 4B**). After adjusting for genotype-by-treatment interaction effects, differential expression analysis revealed 364 DEG unique to WT responses and 944 DEG unique to KO responses. Using these unique transcriptional signatures, Reactome pathway analyses identified pathways associated with WT and KO responses to trauma/HS and FFP resuscitation (**Figure 4C, D**). In WT mice, the top pathways were RNA metabolism, TP53 activation, and cellular responses to stress (**Figure 4C**). Conversely, KO mice that underwent trauma/HS and FFP resuscitation exhibited enrichment of genes associated with pathways related to collagen deposition and integrin cell surface interactions compared to sham mice (**Figure 4D**). Next, we assessed expression levels of select genes involved in pathways known to regulate post-injury organ damage, including thromboinflammation, vascular barrier function, and cell stress responses. Data for WT and KO sham and FFP resuscitated groups were averaged and presented as a heat map (**Figure 4E**). Compared to sham WT mice, sham KO mice exhibited notable baseline reductions in genes that regulate vascular barrier function and cytoprotection. We also noted increased expression of genes that promote inflammation in KO sham versus WT sham mice (**Figure 4E**). When evaluating the differences in responses to trauma/HS by genotype, we found robust overall transcriptomic changes between sham and trauma/HS and FFP resuscitation in WT mice that were diminished in KO mice. While FFP resuscitation reduced expression of genes related to coagulation in WT mice compared to sham, this was not the case with KO mice as they exhibited sustained overall expression of coagulation genes after FFP resuscitation. Similarly, while FFP resuscitation reduced expression of genes related to barrier function in WT mice, minimal changes were observed between KO sham and FFP resuscitated mice. Finally, in WT mice, trauma/HS and FFP resuscitation was associated with increased cytokine gene expression but reductions in endothelial adhesion molecules, such as ICAM and VCAM, and stress response genes, including *Bcl2*, *Txnip*, and *Ddit4*. In contrast, FFP resuscitation in KO mice did not suppress inflammatory gene expression and increased expression of the stress response gene, *Nqo1* (**Figure 4E**).

**Figure 4.**
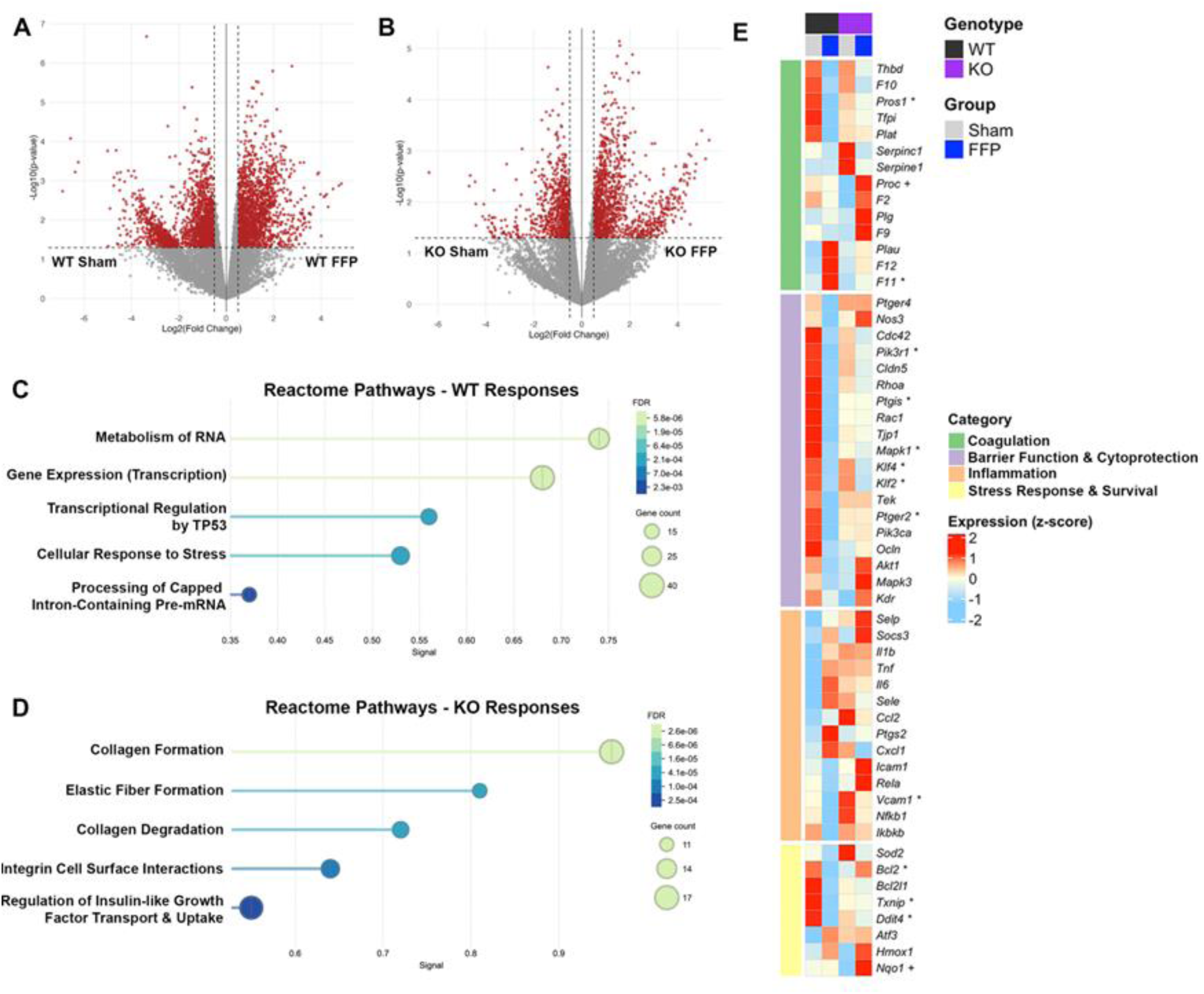
*Hs3st1* KO and WT mice have distinct transcriptomic signatures after trauma and HS. RNA was extracted from pulmonary tissue of sham and FFP-resuscitation WT and KO and subjected to bulk RNAseq analysis. Differential gene expression signatures between sham and FFP (A) WT and (B) KO mice are depicted in volcano plots. The top five Reactome pathways associated with (C) WT and (D) KO responses are shown. (E) Expression levels of genes potentially influenced by AT-3OS HS signaling. N=4-6 mice per group. Significant differences between genes are indicated by * for WT mice and + for KO mice (sham vs FFP, Welch’s t-test, p < 0.05).

### Loss of EC HS3ST1 promotes leukocyte engagement during inflammation in vitro

Given our in vivo findings concerning altered EC activation and leukocyte infiltration, we next performed in vitro perfusion experiments to evaluate if human PMN more readily adhered to HS3ST1 deficient EC, thus demonstrating orthogonal conservation of HS3ST1 function in leukocyte:EC interactions. KO EC were more sensitive to TNFα stimulation (100 ng/mL) as compared to WT EC (**Supplemental Figure 2**), thus a suboptimal dose of TNFα (50 ng/mL) was used to stimulate WT and KO EC to induce PMN rolling and adherence. We found that TNFα-stimulated KO EC had increased frequencies of slow (0-50 µm/sec; p<0.01) and fast rolling PMN (50-250 µm/sec; p<0.01) along their surface and a decrease in the frequency of non-engaging PMN (>250 µm/sec; p<0.0001) as compared to unstimulated KO EC (**Figure 5A**). Of note, these differences in rolling kinetics were more readily observable in KO EC as compared to WT EC. Furthermore, while incubation of WT EC with AT prior to TNFα stimulation led to a reduction in fast rolling PMN (50-250 µm/sec, p<0.0001; **Figure 5B**), AT treatment had no effect on PMN rolling in TNFα-stimulated KO EC (**Figure 5C**).

**Figure 5.**
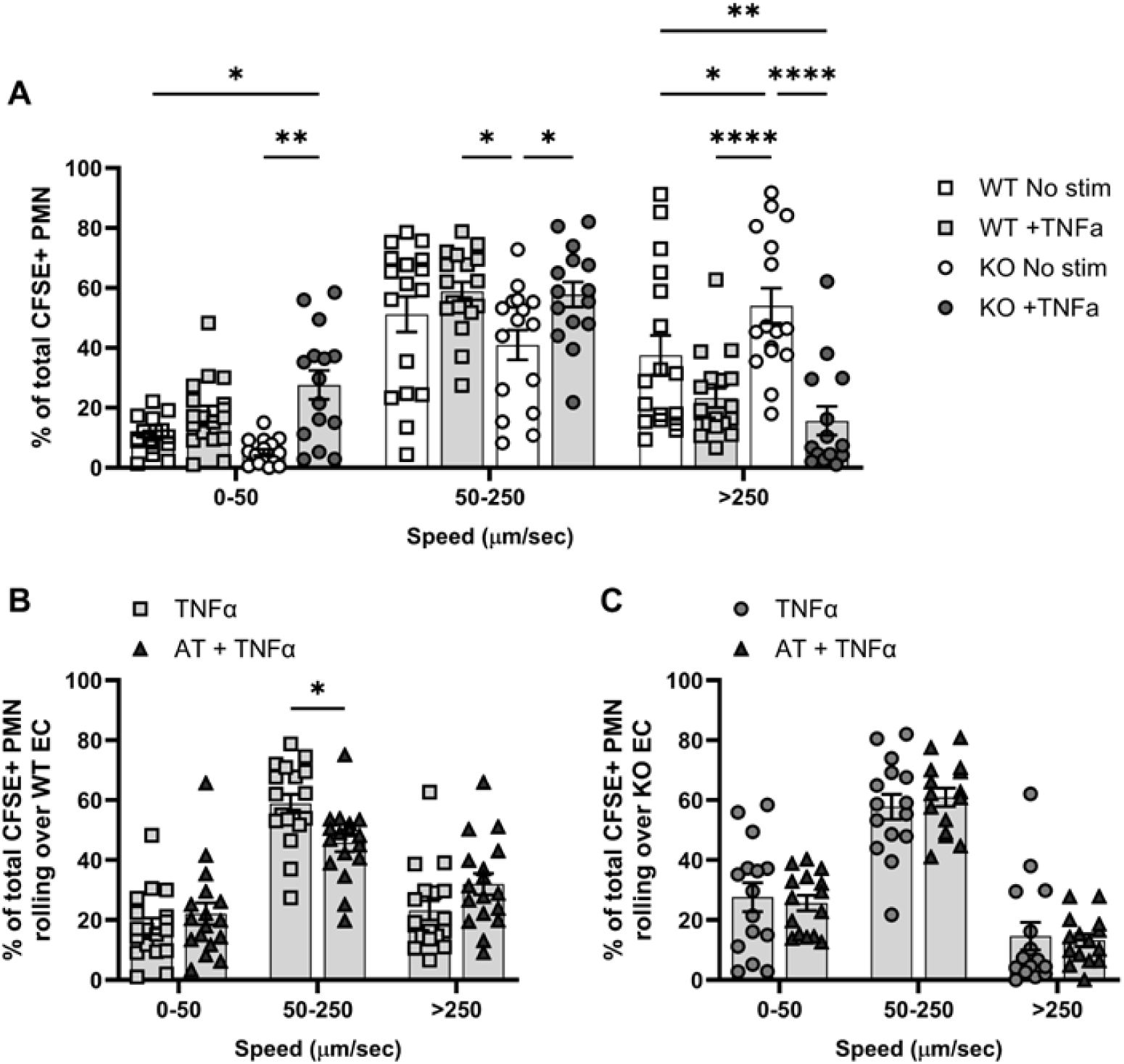
Increased PMN Rolling and Adherence to HS3ST1 Deficient EC after TNFα Stimulation. (A) The frequency of PMN at each speed (0-50 µm/sec, slow rolling and adherent; 50-250 µm/sec, fast rolling; >250 µm/sec, non-engaging) in WT and KO EC in the absence (“No stim”) and presence (“+ TNFα”) of TNFα stimulation (50 ng/mL overnight). N=5-7 unique fields per channel, with one channel per unique EC passage:human PMN donor pairing; thus 15-20 data points per condition. Comparison of the effects of AT (150 µg/mL, 30 min) incubation prior to TNFα stimulation in (B) WT EC and (C) KO EC. Significant differences between EC genotype + stimulation conditions are indicated (two-way ANOVA, with a Tukey’s multiple comparison test across groups within a given speed; *, p < 0.05; **, p < 0.01; ****, p < 0.0001).

### Trauma Patient Demographics and Outcomes

A total of 472 patients met the criteria for inclusion in this analysis and had blood available for DNA extraction. Thirty-one patients (7%) were homozygous for the rs16881446 variant allele (G/G genotype), 190 (40%) were heterozygous (A/G), and 251 (53%) were homozygous for the WT allele (A/A). This is consistent with allele frequencies previously reported in larger cohorts.(9) Patient demographics, injury characteristics, transfusion requirements and outcomes are presented in **Table 1**. When comparing patients across genotypes, we observed no significant differences in patient age, sex, race, or admission vitals between groups. In addition, injury mechanism, severity, and overall transfusions were similar between groups. Ventilator- and ICU-free days were lower in patients with the G/G genotype; however, this did not meet the threshold for significance (p=0.05 and p=0.09, respectively). The incidence of in-hospital mortality was significantly higher among A/G and G/G patients compared to A/A patients (A/A: 15.5% vs A/G: 26.3% vs G/G 25.8%; p=0.02) (**Table 1**).

**TABLE 1.**
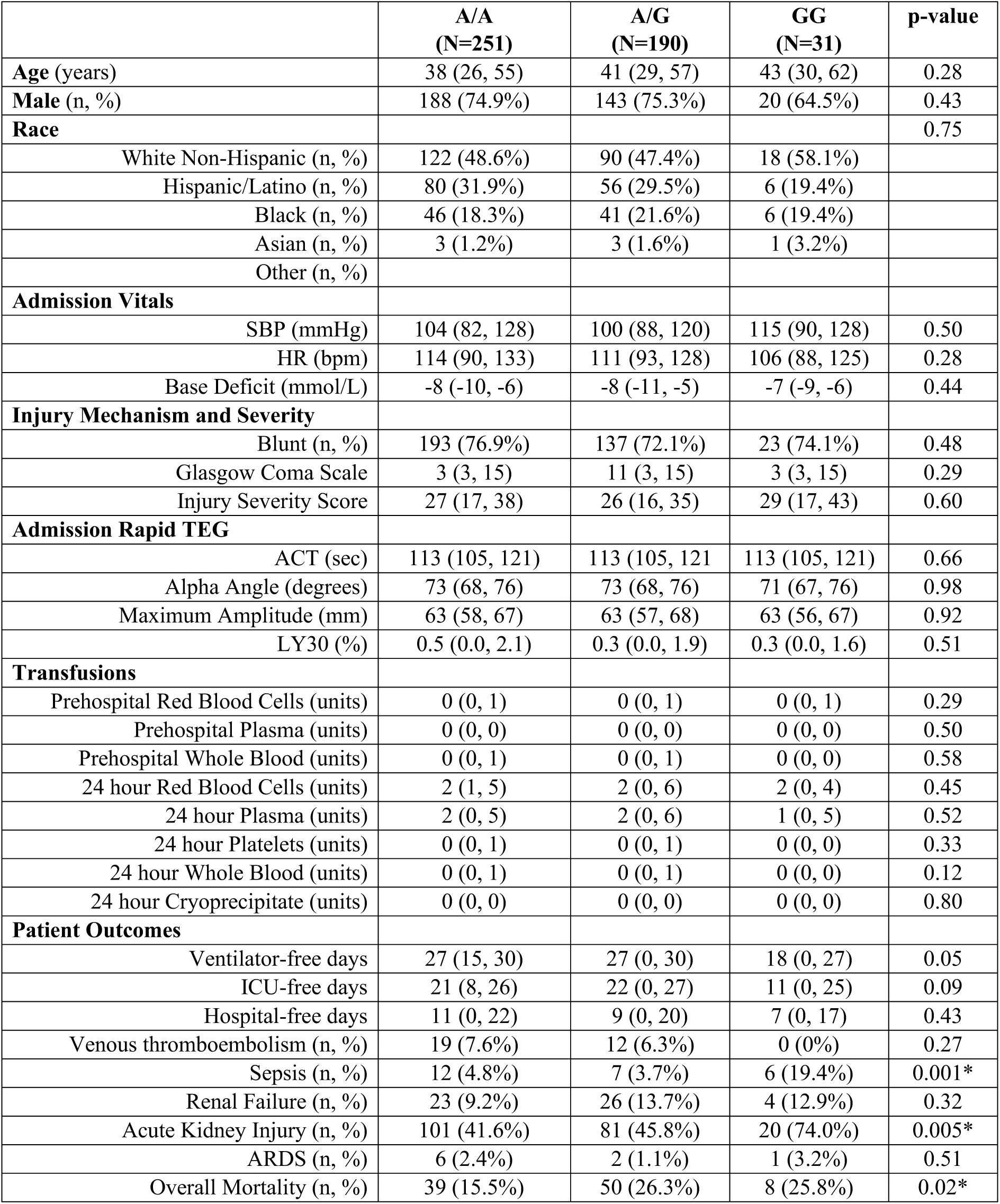
Patient demographics, injury characteristics and outcomes by rs16881446 genotype. Data presented as median (IQR) for continuous variables.

When assessing in-hospital complications, we found that the incidence of sepsis was significantly higher in patients with the G/G genotype (19.4%) compared to A/A (4.8%) or A/G (3.7%) patients (p=0.001) (**Table 1**). In addition, the incidence of acute kidney injury (AKI), defined as KDIGO stage ≥2, was significantly higher in G/G patients (74.0%) compared to A/A (41.6%) or A/G (45.8%) patients (p=0.005) (**Table 1**). No differences in the frequency of VTE, renal failure or acute respiratory distress syndrome diagnoses were identified. There were no differences in admission thromboelastographic parameters between groups, including activated clotting time (ACT), alpha angle, maximum amplitude, and percent lysis at 30 minutes (LY30) (**Table 1**).

### Patients with the HS3ST1 rs16881446 variant polymorphism exhibit increased systemic inflammation, but no change in hypercoagulability

Patients with the A/A genotype were matched with G/G patients based on age, sex, injury mechanism, and injury severity. Plasma inflammatory mediators were measured using a pre-specified Mesoscale Discovery multiplex assay. We found that compared to A/A patients, those with the G/G genotype had significantly increased plasma levels of interleukin (IL)-8 (**Figure 6A**), tumor necrosis factor (TNF)-β (**Figure 6B**), macrophage chemoattractant protein (MCP)-1 (**Figure 6C**), TEK receptor tyrosine kinase (TIE-2) (**Figure 6D**), and significantly reduced levels of IL-17 (**Figure 6E**). Marginal but not significant increases in C-reactive protein were also noted among G/G patients (**Figure 6F**).

**Figure 6.**
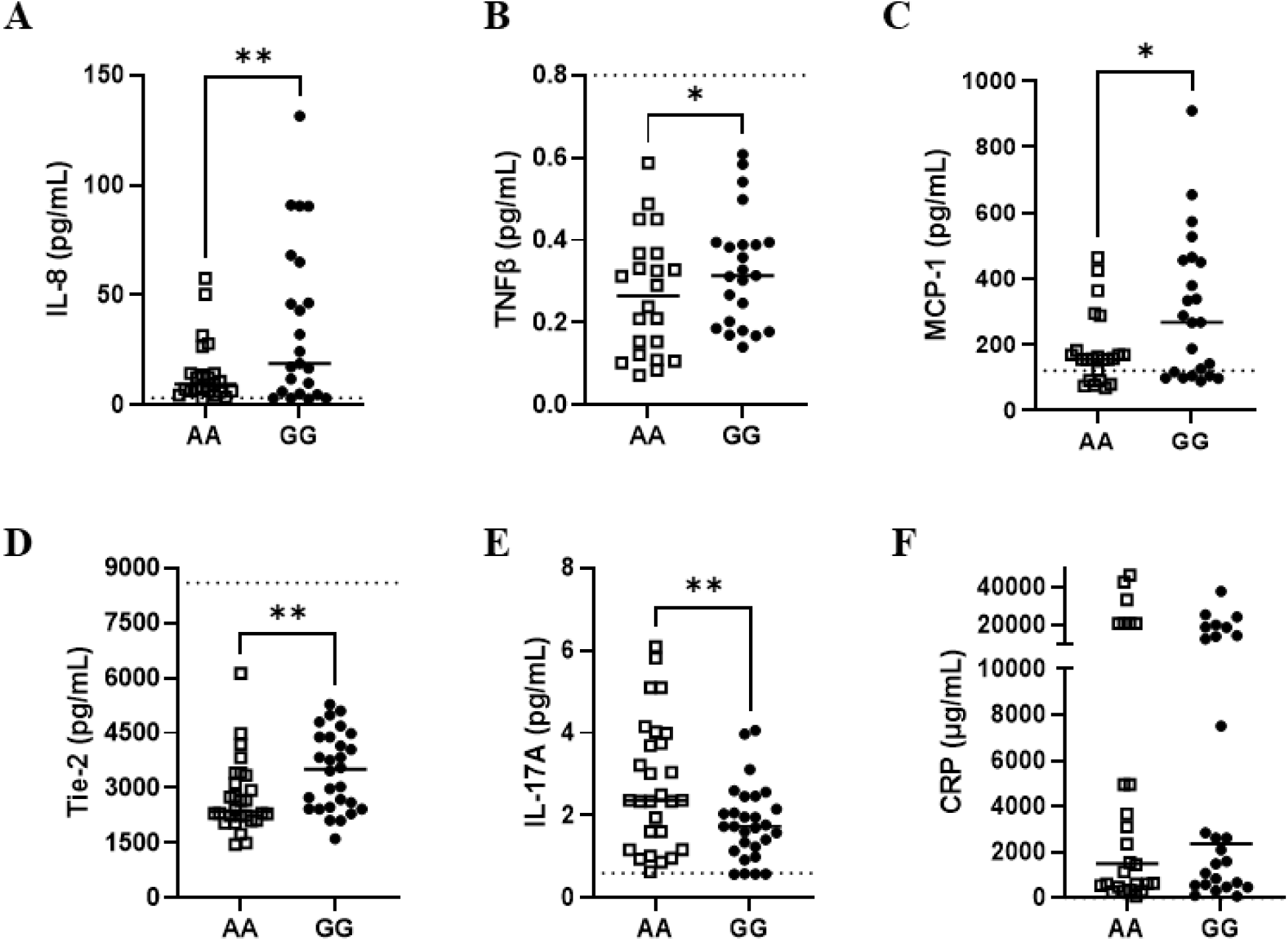
Plasma inflammatory mediators are elevated upon admission in rs16881446 G/G patients compared to A/A patients. Inflammatory mediators were quantified in plasma samples collected upon admission to the emergency room in G/G and A/A patients matched based on age, sex, injury mechanism and severity. Data were Log2 transformed and presented as a heat map (A). Raw values are plotted for markers that were found to significantly differ between groups (B). * denotes p<0.05, Dotted line shows mean levels in healthy subjects, according to manufacturer.

To determine whether rs16881446 modulates circulating levels of AT or changes in activation of coagulation, plasma levels of AT antigen and thrombin-antithrombin (TAT) complex were measured by ELISA. While we observed significant decreases in plasma AT compared to healthy donors, as previously described(20–22), there we no differences between A/A and G/G patients (**Figure 7A**). Moreover, TAT complex levels were significantly elevated in both groups compared to healthy, but no differences were found between A/A and G/G patients (**Figure 7B**).

**Figure 7.**
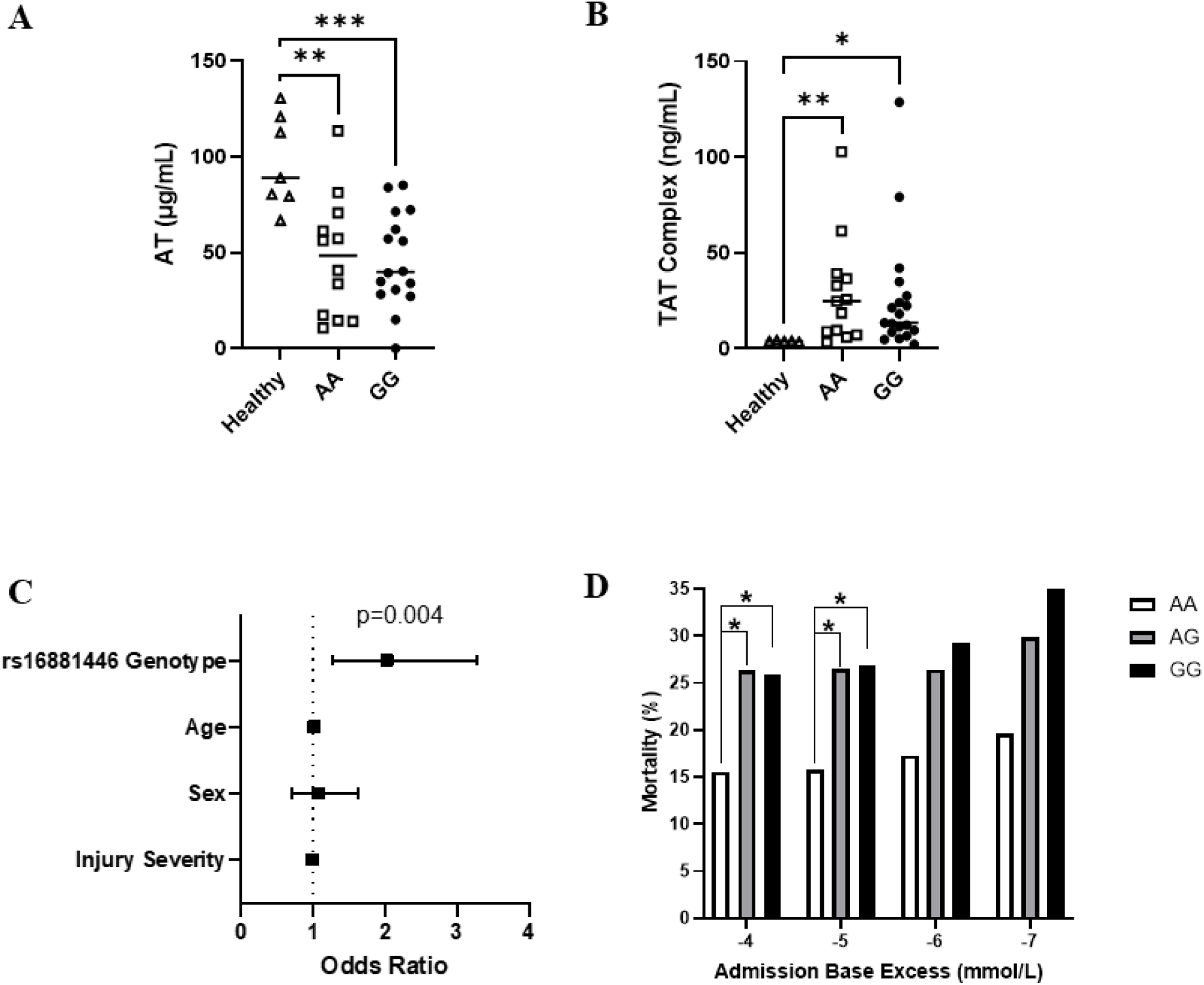
HS3ST1 genotype does not modulate AT levels or hypercoagulablity after trauma but is an independent predictor of mortality. Admission plasma levels of antithrombin (AT) (A) and thrombin-antithrombin complex (TAT) (B) were measured by ELISA in G/G and A/A patients matched based on age, sex, injury mechanism and severity. Multivariable logistic regression analyses were performed to determine the independent contribution of rs16881446 genotype, age, sex, and injury severity to death and odds ratios presented (C). The incidence of mortality by genotype and across admission base excess values are provided (D). * denotes p<0.05

### HS3ST1 rs16881446 Genotype is an Independent Predictor of Mortality after trauma

Multivariable logistic regression analyses were performed to determine the independent association between rs16881446 genotype and mortality (**Table 2**, **Figure 7C**). The rs16881446 genotype was independently associated with an increased risk of death (Odds Ratio 2.03, 95% CI 1.26, 3.27; p=0.004) even when controlling for clinical covariates associated with mortality, including age, sex, and injury severity scores (**Table 2**, **Figure 7C**). When comparing the incidence of death across genotypes with increasing shock severity, we found that mortality was significantly higher in both A/G and G/G patients compared to A/A patients among those with higher admission base excess values (−4 and -5 mmol/L; both p<0.05) (**Figure 7D**); however, as the severity of shock increased, we observed divergence between A/G and G/G patients with a higher incidence of mortality in G/G patients, although due to limitations in sample size this was not statistically significant.

**TABLE 2.**
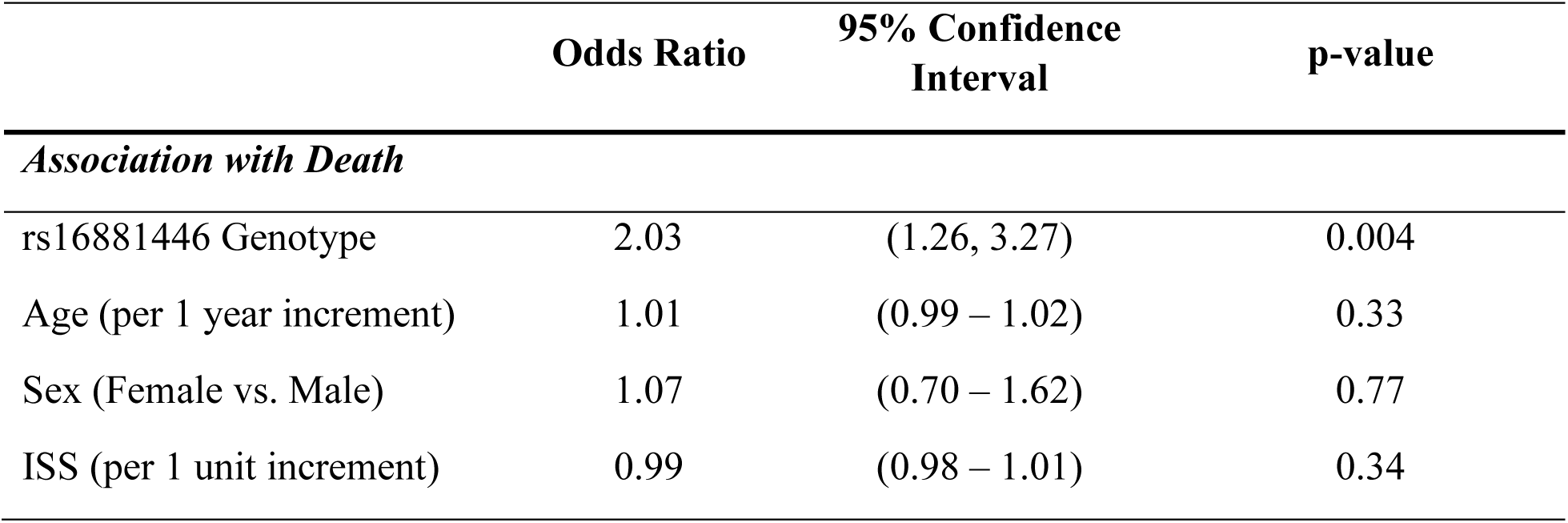
Multivariable logistic regression model evaluating all HS3ST1 rs16881446 genotypes and mortality, while controlling for age, sex, and injury severity score.

|  | **Odds Ratio** | **95% Confidence Interval** | **p-value** |
| --- | --- | --- | --- |
| *Association with Death* |  |  |  |
| rs16881446 Genotype | 2.03 | (1.26, 3.27) | 0.004 |
| Age (per 1 year increment) | 1.01 | (0.99 – 1.02) | 0.33 |
| Sex (Female vs. Male) | 1.07 | (0.70 – 1.62) | 0.77 |
| ISS (per 1 unit increment) | 0.99 | (0.98 – 1.01) | 0.34 |

## Discussion

Over the past 15 years, trauma resuscitation has undergone a revolutionary development characterized by a move towards plasma and whole blood-based resuscitation.(23) These changes initially reduced morbidity and mortality before reaching a plateau in clinical effect. Indeed, gaps in our understanding of the mechanistic underpinnings driving thromboinflammation and post-injury organ damage have precluded development of targeted therapeutics to improve long-term outcomes after trauma. Our prior work has demonstrated that trauma and HS result in modifications to the EC heparan sulfate landscape that resulted in reduced expression of 3-OS heparan sulfate, both in rodents and humans.(15, 24) Here, we show that mice deficient in *Hs3st1*, the gene responsible for EC production of 3-OS heparan sulfate, develop more severe pulmonary inflammation and histologic injury compared to WT controls after trauma and HS, without altering fibrin deposition. Additionally, we found that EC loss of HS3ST1 led to increased human PMN rolling and adherence to inflamed EC, mirroring the data in our mouse model. Moreover, while AT treatment reduced PMN adherence to WT cells, it had no protective effects in HS3ST1 KO cells. We also show that the HS3ST1 loss-of-function rs16881446 variant polymorphism is an independent predictor of mortality in trauma patients. Notably, in both mice and humans we found that HS3ST1 deficiency was associated with increased systemic inflammation, but not hypercoagulability.

Similar to heparin, 3-OS heparan sulfate binds the endogenous anticoagulant AT through a pentasaccharide motif that accelerates AT-mediated inhibition of coagulation proteases, including factor Xa and thrombin, in vitro. It was therefore widely regarded that 3-OS heparan sulfate was a vital mediator of coagulation; however, prior work showed that hemostasis in healthy *Hs3st1* KO mice was normal, despite a 90% reduction in AT binding to blood vessels.(13) Moreover, the propensity for thrombosis was unchanged.(13) This led to speculation that 3-OS heparan sulfate is mainly necessary for anticoagulation during dysregulated, hypercoagulable states and/or its role in immunomodulation predominates in vivo. Indeed, when exposed to endotoxemic challenge, *Hs3st1* KO mice exhibited accelerated lethality, in part, due to a profound sensitivity to TNFα and were resistant to the therapeutic and anti-inflammatory properties of AT administration.(9) Our data are in agreement with this prior work, showing that compared to WT, *Hs3st1* KO mice subjected to trauma and HS develop greater pulmonary inflammation and more severe histologic lung injury after resuscitation with LR, but no differences in local fibrin deposition. Notably, while resuscitation with FFP, which contains AT, reduced lung inflammation and injury in WT mice, *Hs3st1* KO mice were resistant to the organ protective effects of FFP, displaying similar lung damage to those of mice resuscitated with LR. This was not due to enhanced susceptibility or altered reversal of HS, as metabolic indicators of shock and resuscitation volumes were similar across groups. These findings mirror those of our prior work using a heparan sulfate antagonist, surfen, to block AT-EC interactions.^15^ Moreover, these same observations were found in our in vitro perfusion system assessing human PMN adherence to EC: loss of HS3ST1 led to greater PMN rolling and engagement after TNF stimulation of EC, and that supplementation with AT was anti-inflammatory in WT EC but had no effect on KO EC. Thus, the role of HS3ST1 in modulating leukocyte recruitment and engagement dynamics under flow appears to be conserved across species.

Using RNAseq to examine lung transcriptomic signatures, we found that compared to sham WT mice, sham *Hs3st1* KO mice have reduced expression of genes that promote vascular barrier integrity and increased expression of genes that control innate immune responses. However, baseline lung histology and presence of leukocyte infiltrates were similar. This indicates that 3-OS heparan sulfate is important for maintenance of anti-inflammatory homeostasis during health and *Hs3st1* KO mice are transcriptionally primed to respond to acute insults, resulting in enhanced organ inflammation. Interestingly, we found that while trauma/HS and FFP resuscitation modulated overall gene expression patterns, including reductions in genes that promote coagulation, barrier function, endothelial-leukocyte adhesion, and cell stress, this was not the case in KO mice. Here, we found that KO mice had an attenuated transcriptional response to FFP, with minimal suppression of genes that promote thromboinflammation. In addition, we observed increased *Proc* expression, which regulates production of protein C, in FFP KO mice compared to sham, which could indicate compensatory anticoagulant responses induced when 3-OS heparan sulfate is absent. While these analyses are intended to be exploratory and hypothesis generating, they reinforce the premise that *Hs3st1* KO mice exhibit basal vascular permeability and inflammation, implicating 3-OS heparan sulfate and AT-mediated anti-inflammatory functions with protection against post-trauma organ injury.

To extend these findings translationally, we examined the link between trauma patient clinical outcomes and the *HS3ST1* rs16881446 polymorphism, a gene mutation in the putative transcriptional regulatory region that significantly reduces endothelial expression of HS3ST1.(9) The frequency of homozygosity for the variant allele (G/G genotype) was 7% in this population. While there was no difference in demographics, injury mechanism/severity, or transfusion treatment between the genotypes, we found that G/G patients had a significantly higher incidence of developing sepsis and acute kidney injury; moreover, mortality was higher among patients with either the A/G or G/G genotypes. This indicates that having even a single variant allele increases the risk of death after trauma and HS. In agreement with this, we found that rs16881446 genotype status (A/G or G/G vs. A/A) was an independent predictor of mortality, even when controlled for age, sex, and injury severity. Interestingly, we further found that compared to age, sex, and injury matched A/A patients, those with the G/G genotype exhibited significantly increased systemic inflammatory markers upon admission, but no change in markers of hypercoagulability. In keeping with our results, prior work from Smits et al. demonstrated that among patients undergoing coronary catheterization, the G/G genotype was associated with both more severe coronary artery disease and a higher incidence of experiencing heart attack, coronary intervention or bypass graft, transient ischemic attack, stroke, or occlusive vascular disease.(9) In addition, recent work from Calusen et al. showed that suppression of HS3ST1 during pancreatic tumor progression promotes inflammation in the tumor microenvironment, promoting metastasis.(25) Together, these studies suggest that 3-OS heparan sulfate is an important regulator of inflammation, a function that predominates in vivo over its anticoagulant function, and a key determinant of outcomes during acute illness, chronic cardiovascular disease, and cancer. In addition, rapid genotyping of rs16881446 could offer a novel screening tool for identifying patients at highest risk of post-trauma complications, despite adequate blood product resuscitation.

The findings herein suggest that targeting 3-OS heparan sulfate could be a viable, directed therapeutic option for pharmaceutical development to improve outcomes after trauma, particularly among patients with the G/G genotype. Recent advances in the chemoenzymatic methods have enabled production of synthetic, structurally homogenous, and reversable heparan sulfate molecules with precisely controlled sulfation patterning.(26–30) Previous studies have established the potential therapeutic benefits of administering synthetic heparan sulfates for liver damage associated with acetaminophen,(31) ischemia reperfusion injury,(32) and sepsis.(33) Recently, our group has demonstrated that synthetic 3-OS heparan sulfate, a 12-mer oligosaccharide, significantly attenuates multi-organ injury following trauma and HS in rats, without compromising hemodynamic stability or hemostatic function.(17) Future studies leveraging large animal models will determine whether this approach is safe and potentially effective for use in humans and testing in clinical trials.

This study has several limitations. Due to intrauterine growth retardation associated with breeding *Hs3st1* heterozygotes, WT and KO mice are smaller than commercial C57BL/6 mice, making collection of plasma for analysis challenging after HS and limiting our ability to compare systemic thromboinflammatory markers, as we did in patients. In addition, while the 7% frequency of rs16881446 G/G is moderately high among polymorphisms, our limited patient cohort size may have reduced our ability to detect meaningful differences in clinical outcomes or plasma markers between groups. Larger, multicenter studies should be conducted to validate the prognostic value of rs16881446 genotyping and increase generalizability across institutions. Future studies evaluating the role of these SNPs in human EC biology and the effect on coagulation and leukocyte recruitment will be useful for further defining the mechanisms surrounding increased mortality in trauma patients.

## Conclusions

In conclusion, *Hs3st1* deficiency is a key driver of lung inflammation and injury after trauma and HS. EC-specific loss of HS3ST1 resulted in increased leukocyte adherence under flow in the setting of TNFα-induced activation. Lastly, the loss-of-function SNP in the *HS3ST1* gene, rs16881446, is independently associated with poor outcomes in trauma and HS patients. These findings suggest that 3-OS heparan sulfate is an important regulator of post-trauma inflammation and organ protection during critical illness that could be targeted for improved risk stratification and directed therapies to improve outcomes after severe injury and HS.

## Acknowledgements

This project was supported by awards from the National Institutes of Health, including R35GM146859 to JCC, R35GM137958 to JRR, R01HL171594 to KAT, and R01HL182062 to MJC.

## Author Contributions

AM designed the research, conducted experiments, analyzed data, and wrote the manuscript. MEC conducted experiments, analyzed data, and edited the manuscript. JK conducted experiments, analyzed data, and edited the manuscript. KAT secured funding, designed, analyzed, and interpreted the perfusion experiments, as well as edited the manuscript. AC designed, performed, and analyzed the perfusion experiments. MP conducted experiments, analyzed data, and edited the manuscript. BO performed experiments and edited the manuscript. MV performed experiments and edited the manuscript. ARM analyzed data and edited the manuscript. CW assisted with study design, oversaw patient sample collection and edited the manuscript. Y-WW performed experiments and edited the manuscript. DJO scored histopathology, analyzed data and edited the manuscript. MJC assisted with data analysis, interpretation, and edited the manuscript. JR assisted with data analysis, interpretation, and edited the manuscript. NS designed the research, interpreted the data, and edited the manuscript. JC conceived the project, secured funding, designed the research, directed the project, analyzed and interpreted data, and wrote the manuscript. All authors contributed to and approved the manuscript.

---

**Supplemental Figure 1.**
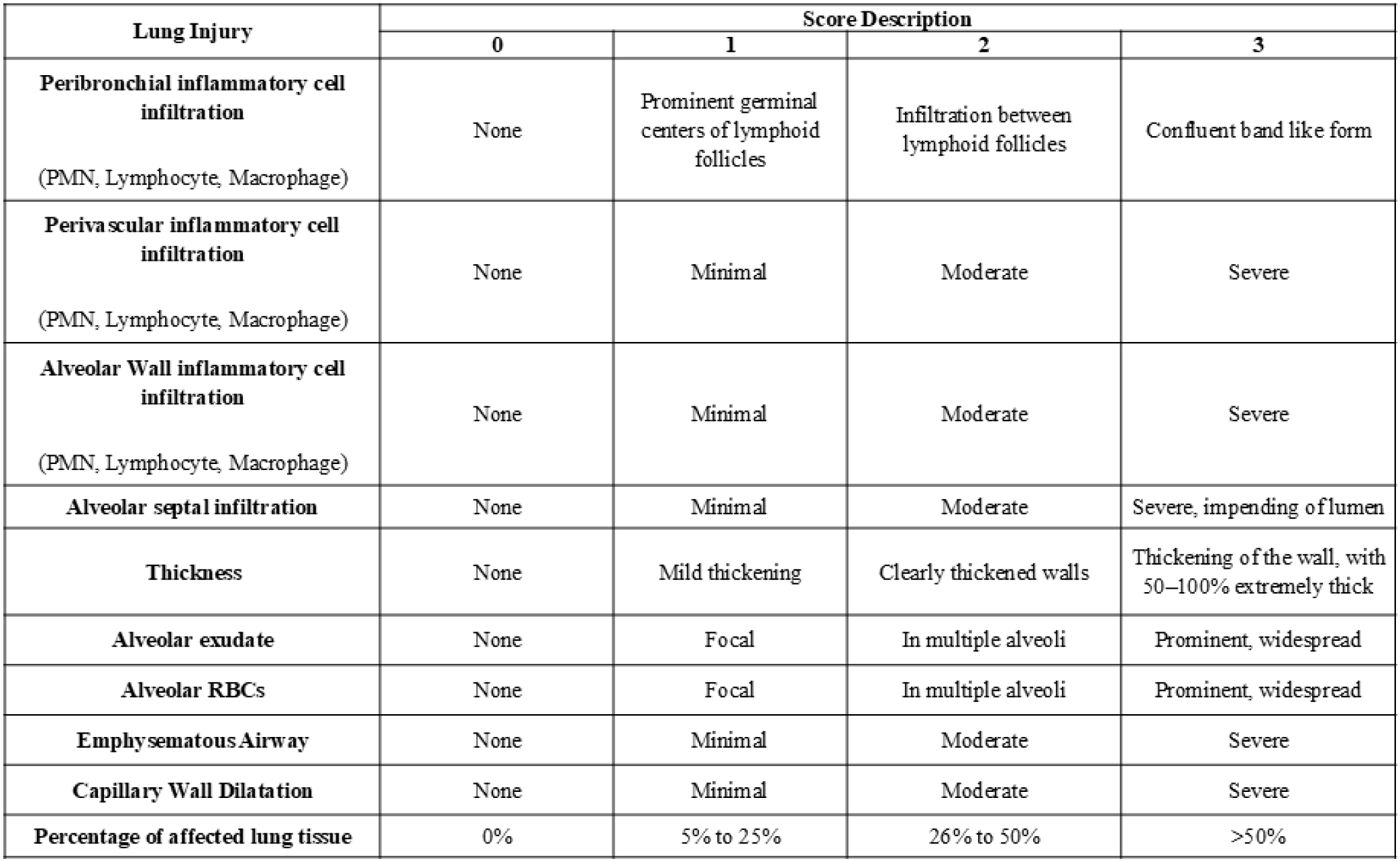
Histologic lung injury scoring scale.

**Supplemental Figure 2.**
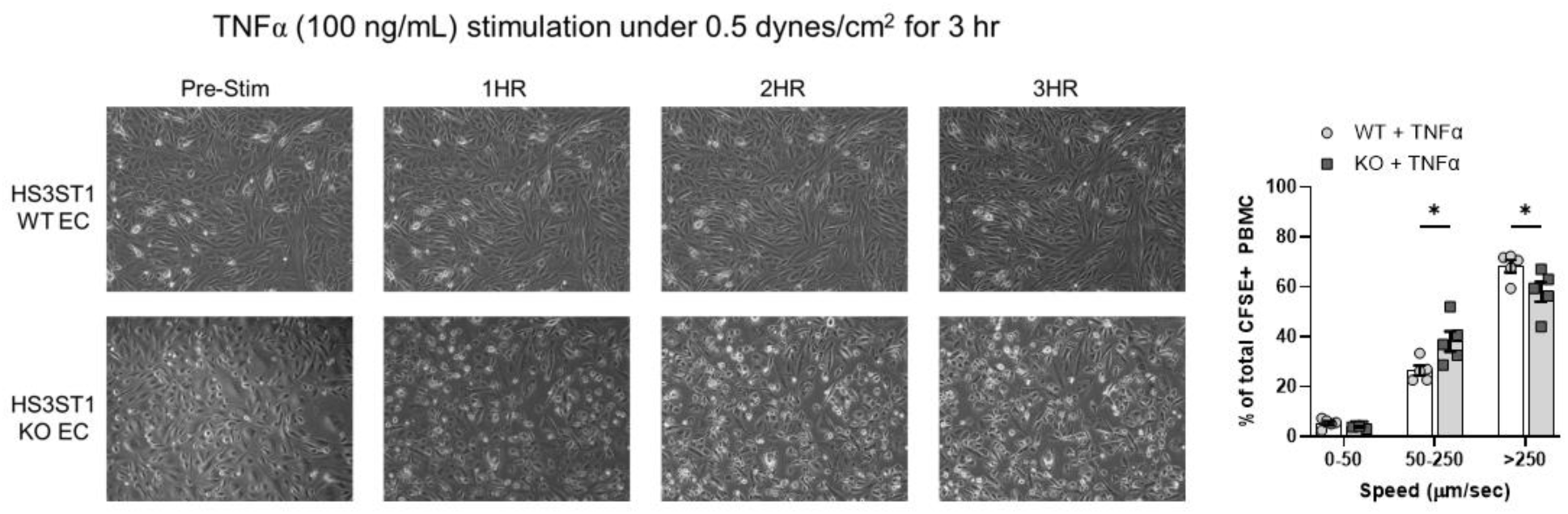
HS3ST1 WT and KO cells are more sensitive to TNFα stimulation. The frequency of PMN at each speed (0-50 µm/sec, slow rolling and adherent; 50-250 µm/sec, fast rolling; >250 µm/sec, non-engaging) in WT and KO EC in the absence (“No stim”) and presence (“+ TNFα”) of TNFα stimulation (100 ng/mL overnight). N=5-7 unique fields per channel, with one channel per unique EC passage:human PMN donor pairing; thus 15-20 data points per condition.

**Supplemental Videos 1 & 2**

Twenty second videos of CFSE labeled PMN rolling over either unstimulated WT EC (Supplemental Video 1) or WT EC stimulated with TNFα (50 ng/mL, overnight) (Supplemental Video 2). The GFP signal from these videos is used in conjunction with NIS Elements AR software to enumerate ethe number and speed of PMN engaging with EC.

## Notes

### Competing Interest Statement

The authors have declared no competing interest.

