## Supplemental Data file for "HS3ST1 regulates pulmonary inflammation and is a determinant of clinical outcomes after trauma and hemorrhagic shock"

**Supplemental Figure 1. Histologic lung injury scoring scale.**

| Lung Injury | Score Description | | | |
|---|---|---|---|---|
| | 0 | 1 | 2 | 3 |
| **Peribronchial inflammatory cell infiltration** (PMN, Lymphocyte, Macrophage) | None | Prominent germinal centers of lymphoid follicles | Infiltration between lymphoid follicles | Confluent band like form |
| **Perivascular inflammatory cell infiltration** (PMN, Lymphocyte, Macrophage) | None | Minimal | Moderate | Severe |
| **Alveolar Wall inflammatory cell infiltration** (PMN, Lymphocyte, Macrophage) | None | Minimal | Moderate | Severe |
| **Alveolar septal infiltration** | None | Minimal | Moderate | Severe, impending of lumen |
| **Thickness** | None | Mild thickening | Clearly thickened walls | Thickening of the wall, with 50–100% extremely thick |
| **Alveolar exudate** | None | Focal | In multiple alveoli | Prominent, widespread |
| **Alveolar RBCs** | None | Focal | In multiple alveoli | Prominent, widespread |
| **Emphysematous Airway** | None | Minimal | Moderate | Severe |
| **Capillary Wall Dilatation** | None | Minimal | Moderate | Severe |
| **Percentage of affected lung tissue** | 0% | 5% to 25% | 26% to 50% | >50% |

**Supplemental Figure 2. HS3ST1 WT and KO cells are more sensitive to TNFα stimulation.** The frequency of PMN at each speed (0-50 µm/sec, slow rolling and adherent; 50-250 µm/sec, fast rolling; >250 µm/sec, non-engaging) in WT and KO EC in the absence ("No stim") and presence ("+ TNFα") of TNFα stimulation (100 ng/mL overnight). N=5-7 unique fields per channel, with one channel per unique EC passage:human PMN donor pairing; thus 15-20 data points per condition.

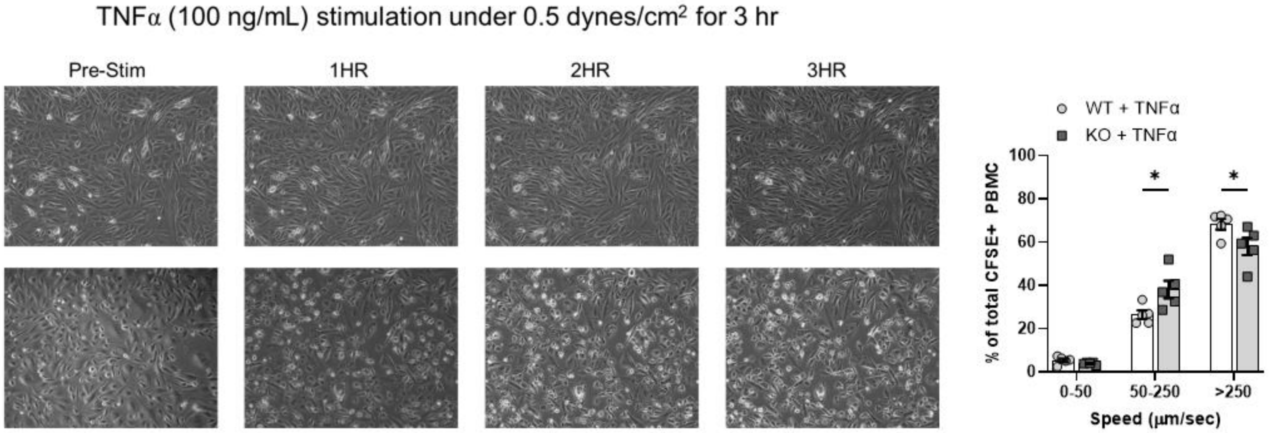

**Supplemental Videos 1 & 2**

Twenty second videos of CFSE labeled PMN rolling over either unstimulated WT EC (Supplemental Video 1) or WT EC stimulated with TNFα (50 ng/mL, overnight) (Supplemental Video 2). The GFP signal from these videos is used in conjunction with NIS Elements AR software to enumerate ethe number and speed of PMN engaging with EC.
